# Cerebellar granule cell migration and folia development requires Mllt11/Af1q/Tcf7c

**DOI:** 10.1101/2023.09.12.557458

**Authors:** Marley Blommers, Danielle Stanton-Turcotte, Emily A. Witt, Mohsen Heidari, Angelo Iulianella

## Abstract

The organization of neurons into distinct layers, known as lamination, is a common feature of the nervous system. This process, which arises from the direct coupling of neurogenesis and neuronal migration, plays a crucial role in the development of the cerebellum, a structure exhibiting a distinct folding cytoarchitecture with cells arranged in discrete layers. Disruptions to neuronal migration can lead to various neurodevelopmental disorders, highlighting the significance of understanding the molecular regulation of lamination. Here we report a role Mllt11/Af1q/Tcf7c (Myeloid/lymphoid or mixed-lineage leukemia; translocated to chromosome 11/All1 Fused Gene From Chromosome 1q, also known as Mllt11 transcriptional cofactor 7; henceforth referred to Mllt11) in the migration of cerebellar granule cells (GCs). We now show that Mllt11 plays a role in both tangential and radial migration of GCs. Loss of *Mllt11* led to an accumulation of GC precursors in the Rhombic Lip region and a reduction in the number of GCs successfully populating developing folia. Consequently, this results in smaller folia and an overall reduction in cerebellar size. Furthermore, analysis of the anchoring centers reveals disruptions in the perinatal folia cytoarchitecture, including alterations in the Bergmann glia fiber orientation and reduced infolding of the Purkinje cell plate. Lastly, we demonstrate that Mllt11 interacts with Non-muscle Myosin IIB (NMIIB) and regulates its expression. We propose the dysregulation of NMIIB underlies altered GC migratory behavior. Taken together, the findings reported herein demonstrate a role for Mllt11 in regulating neuronal migration within the developing cerebellum, which is necessary for its proper neuroanatomical organization.

## INTRODUCTION

Neurons navigate complex paths and settle into their proper positions within the central nervous system (CNS) via two general modes of migration, tangential and radial. Radial migration primarily occurs as newly born granule cells (GCs) migrate inward, perpendicular to the pial surface, along a radial glial (RG) fiber scaffold to populate the core (Ballif et al., 2004). Tangential migration precedes radial migration and is characterized by neurons moving parallel to the brain surface, utilizing substrates other than RG, such as diffusible and membrane-bound molecules like Slits and Semaphorins (Metin, Vallee, Rakic, & Bhide, 2008; Tran, Kolodkin, & Bharadwaj, 2007). Both modes of migration are crucial for the development of the mammalian CNS, as they allow neurons from different proliferative zones, with distinct lineages and genetic programs, to come together and form synaptic connections (Hatten, 1993; Marin, Valiente, Ge, & Tsai, 2010). Through a tightly co-regulated process of neurogenesis and migration, a stratified structure emerges as cells are distributed at their site of integration sequentially over developmental time. This process, known as lamination, is a common feature in many CNS structures, such as the neocortex, cerebellum, and retina, and it facilitates neuronal organization into distinct projection circuits (Blommers, Stanton-Turcotte, & Iulianella, 2023; Guy & Staiger, 2017; Stanton-Turcotte et al., 2022).

Disruption of laminar organization in the developing CNS has been implicated in disorders such as lissencephaly, periventricular nodular heterotopia, and subcortical band heterotopia due to mispositioned neurons and altered connections (Liu, 2011). Similarly, defects in neuronal migration have been associated with various disorders, including brain malformation, intellectual disability, psychiatric disease, and epilepsy (Liu, 2011). While these disorders can be linked to more than a single gene disruption, many identified mutations involve genes that encode proteins that either interact with or directly regulate the cytoskeleton (Lasser, Tiber, & Lowery, 2018). Microtubules and actin, key components of the cytoskeleton, are highly enriched in the leading processes of migrating granule cells and regulate cerebellar morphogenesis (Gregory, Edmondson, Hatten, & Mason, 1988; Hatten, 1993; Rivas & Hatten, 1995). Disruptions in actin and microtubules or their associated motor proteins can inhibit the cellular migration (Bellion, Baudoin, Alvarez, Bornens, & Metin, 2005; Kawauchi & Hoshino, 2008; Rivas & Hatten, 1995; Trivedi & Solecki, 2011), and pharmacological inhibition of actin polymerization using cytochalasin B is sufficient to inhibit granule cell migration (Hatten, 1993).

While it is well understood that the cytoskeleton plays a central role in orchestrating cell motility in development and cancer, our knowledge of the potential regulators of this process remains incomplete. To that end we identified a protein called Myeloid/lymphoid or mixed-lineage leukemia; translocated to chromosome 11 or ALL1 fused from chromosome 1q or Mllt11 transcription factor 7 cofactor (Mllt11/Af1q/Tcf7c; hereafter referred to as Mllt11) as a cytoskeletal-interacting protein that is crucially required for neuronal migration in the mammalian cortex and retina (Blommers et al., 2023; Stanton-Turcotte et al., 2022). Mllt11 was initially identified as an oncogene in acute myeloid leukemia caused by translocations, but we discovered that it is expressed in post-mitotic neurons throughout the developing CNS (Stanton-Turcotte et al., 2022; W. Tse, Zhu, Chen, & Cohen, 1995; Yamada, Clark, & Iulianella, 2014). The conditional knockout of *Mllt11* from the superficial cortical projection neurons disrupts their proper radial migration and neuritogenesis and leads to reduction in cortical white matter tracts and projections across the corpus callosum. Furthermore, proteomic analysis in fetal brain lysates revealed Mllt11 interactions with actin, tubulin isoforms, and non-muscle myosins (Stanton-Turcotte et al., 2022). Furthermore, we recently showed that *Mllt11* is required for the migration of retinal ganglion cells and displaced amacrine cells in the retina (Blommers et al., 2023). Although the role of *Mllt11* has been characterized during the formation of the mammalian cortex and retina (Blommers et al., 2023; Stanton-Turcotte et al., 2022; Yamada et al., 2014), we do not know if Mllt11 is also utilized as a regulator of migration in the cerebellum, whose complex formation entails a switch from tangential to radial migration of granule cell precursors. Therefore, we sought to investigate the role of Mllt11 in the neuronal migration during cerebellar morphogenesis.

As a highly laminated neural structure, the cerebellum provides an excellent developmental model for studying the mechanisms of neuronal migration (Fig.1A-D) due to its relatively simple layered cytoarchitecture, consisting of four principal fissures (Fig. 1E) and five cardinal lobes (Fig. 1F). Cerebellar granule cells (GCs), the most numerous type of neuron, undergo expansive proliferative divisions and tangential and radial migration to generate the foliated cytoarchitecture of the amniote cerebellum (Iulianella, Wingate, Moens, & Capaldo, 2019; Rahimi-Balaei, Bergen, Kong, & Marzban, 2018; Xu et al., 2013). Originating in the rhombic lip, GCs must first undergo tangential migration to form the external granular layer, at which point they begin to migrate radially to inhabit the cerebellar core (Rahimi-Balaei et al., 2018). Here we report that the conditional ablation of *Mllt11* from the rhombic lip and external granular layer during cerebellar development alters both the tangential displacement of GC precursors, and inward radial migration of GCs, impacting folia formation. We also report that Mllt11 interacts with Non-muscle Myosin IIB (NMIIB), a key regulator of neuronal migration (Ma, Bao, & Adelstein, 2007; Ma, Kawamoto, Hara, & Adelstein, 2004), whose levels are increased in *Mllt11* mutant brains. We link these observations by proposing a role for Mllt11 in GC migration in part through its interaction with NMIIB.

**Figure 1.**
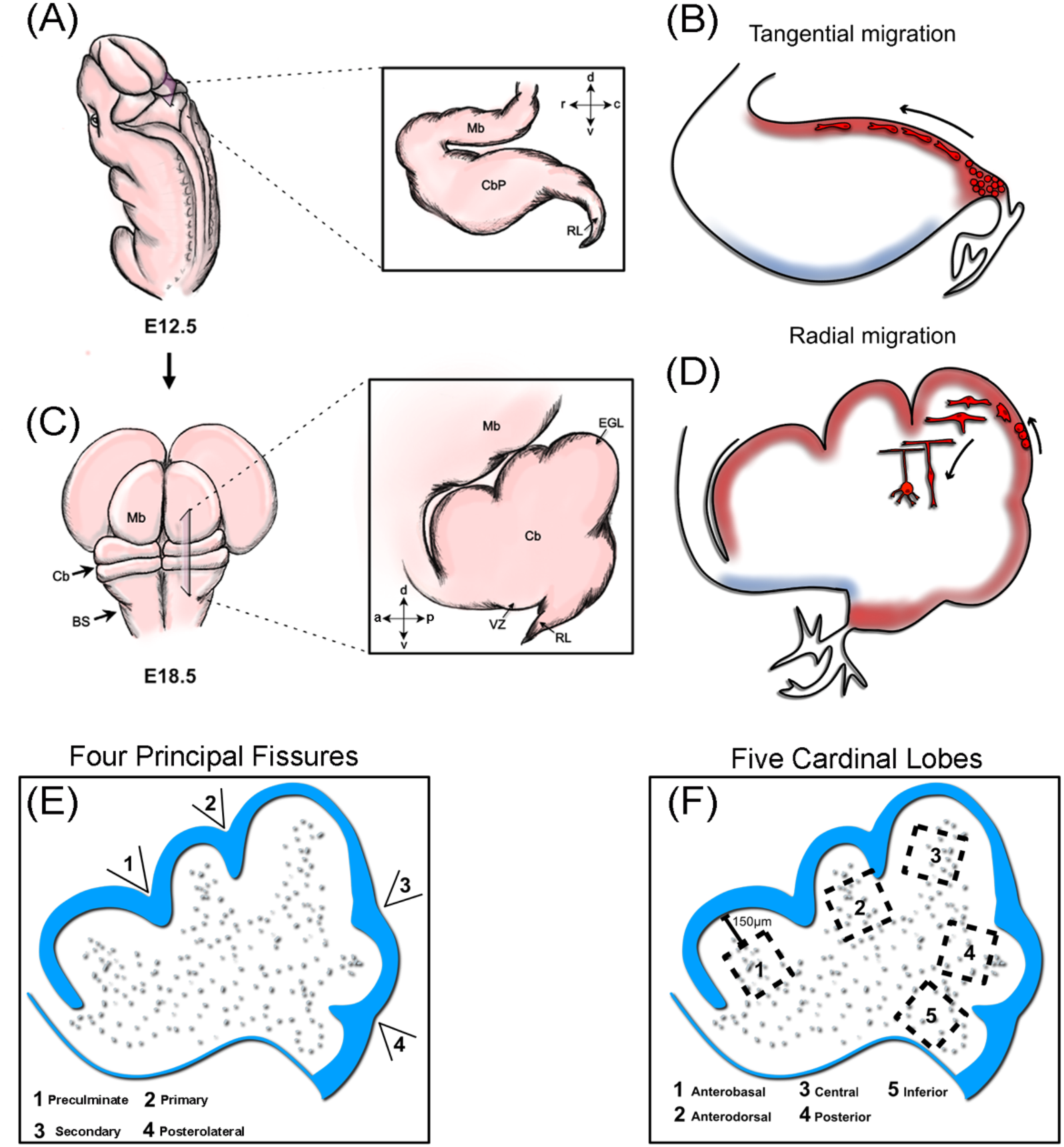
Illustration of cerebellar granular cell migration. **(A)** E12.5 cerebellum posterior view and inset with parasagittal cross-section. At this stage, the cerebellar primordium surface is smooth. **(B)** The first GCPs are born in the RL around E12 and migrate tangentially over the dorsal surface of the cerebellar anlage to form the EGL at E14.5. **(C)** E18.5 cerebellum posterior view and inset with parasagittal cross-section. By this time, four invaginations, or principal fissures, can be identified on its surface which separate as five cardinal lobes. **(D)** Following terminal division, postmitotic GCs move into the inner EGL and emit an axon at each pole in the medial-lateral direction which will become parallel fibers. GCs will undergo nuclear translocation along one parallel fiber before extending a radial process and descending along Bergmann glia fibers into the cerebellar core indicative of a switch to radial migration at E18.5. **(E)** Schematic identifying the four principal fissures of the cerebellum **(F)** Schematic identifying the five cardinal lobes. Abbreviations: BS, brain stem; Cb, Cerebellum; Mb, midbrain; RL, rhombic lip.

## METHODS

### Animals

All animal experiments were done according to approved protocols from the IACUC at Dalhousie University. Mice (*Mus musculus*) carrying a mutation in which the *Mllt11* locus is flanked by loxP sites (floxed) were generated as previously described (Stanton-Turcotte et al., 2022). The Ai9 *Rosa26r^TdTomato^* reporter mouse [B6.Cg-Gt(ROSA)26Sortm9(CAG-tdTomato Hze/J, 007909), The Jackson Laboratory] was crossed into the *Mllt11^flox/flox^* strain, giving rise to *Rosa26r^TdTomato/TdTomato^; Mllt11^flox/flox^* mice and allowing for visualization of cells that had undergone Cre-mediated excision of a floxed translational stop sequence engineered upstream of the *TdTomato* cDNA within the ubiquitously expressed *Rosa26* locus. To delete *Mllt11* within GC of the developing cerebellum, *Cux2^CreERT2/+^; Rosa26r^TdTomato/TdTomato^* mice were crossed with the *Mllt11* floxed mice to generate *Cux2^CreERT2/+^; Rosa26r^TdTomato/TdTomato^*control and *Cux2^CreERT2/+^; Rosa26r^TdTomato/TdTomato^; Mllt11^flox/flox^* conditional knockout (cKO) strains, allowing for excision of *Mllt11* and visualization of recombination events via TdTomato fluorescence in Cux2-expressing cells following tamoxifen-induced Cre activation. 100μL of Tamoxifen (20mg/mL) was administered by intraperitoneal injection at E12.5. Confirmation of *Mllt11* expression in the developing cerebellum was done using the targeted pre-floxed *Mllt11* allele, which houses a *lacZ* cassette encoding β-galactosidase (β-gal; *Mllt11^tm1a(KOMP)Mbp^*). Genotyping was performed following procedures previously described (Stanton-Turcotte et al., 2022). The *Cux2^iresCre^* conditional *Mllt11* knockout used for western blot analysis was described in (Stanton-Turcotte et al., 2022).

### Histology

To harvest the brains of embryonic mice at consistent developmental timepoints, timed mating of mice was conducted. Following a mating pairing female mice were examined daily for presence of vaginal plugs, and embryos were considered to be 0.5 days post conception (E0.5) at noon of the day on which vaginal plugs were identified. Pregnant dams were sacrificed by cervical dislocation and litters were harvested at E12.5, E14.5, E16.5, and E18.5. Embryonic brains were dissected out and fixed in 4% PFA in PB for 2-8 hours depending on developmental stage. Post-fixation, tissue was washed in PBS for 10 minutes and then cryoprotected in 15% and then 30% sucrose. Once equilibrated, whole brains were then embedded in optimum cutting temperature compound (OCT; Tissue-Tek, Torrance, CA) and stored at -80°C until sectioning.

### Immunohistochemistry

Tissue was cryosectioned sagitally at 14μm using a Leica CM1850 cryostat with serial sections placed onto superfrost plus slides (VWR, Radnor, PA). Ten sections were mounted per slide to capture multiple axial levels on each slide. At least three animals were sectioned and analyzed for each stage. Sections were stored at -20°C until stained. Sections were first permeabilized for 10 minutes in PBS + 0.5% TritonX100 (PBT) and subsequently blocked in 3% Donkey Serum in 0.1% PBT for one hour at room temperature (RT). Sections were then incubated with primary antibodies in blocking buffer overnight at 4°C, extensively washed with PBS, and then incubated with secondary antibodies in blocking buffer at RT. Secondary antibodies were rinsed with PBS and nuclei were labeled using DAPI (4’6-diamidino-2-phenylindole; Sigma, St. Louis, MS) in PBS (1:10,000 of 3mg/mL DAPI stock: PBS) by incubation for two minutes at RT. Final PBS rises (3 x 5 minutes) followed and slides were mounted with coverslips and Dako Fluorescent Mounting Medium. The following primary antibodies were used: rabbit anti-Pax6 (1:500, Abcam), rabbit anti-Calbindin D28K (1:500, Sigma), and goat anti-Nestin (1:250, Santa Cruz). Species-specific AlexaFluor 488-, 568-, and 647-conjugated secondary antibodies (Invitrogen/ThermoFisher) were used at 1:1500 to detect primary antibodies.

### *In Situ* Hybridization

In situ hybridization (ISH) was performed on 30μm frozen parasagittal sections obtained from E18.5 control brains fixed overnight as previously described to visualize Mllt11 mRNA in the cerebellum using an *Mllt11* riboprobe (Yamada et al., 2014).

### β-Galactosidase Staining

*Mllt11* locus activity was visualized using a β-gal Tissue Stain kit (MilliporeSigma). Embryos (E12.5 and E14.5) or isolated whole brains (E16.5 and E18.5) were fixed for 30 minutes in fresh 4% PFA in PB, followed by 3 x 10-minute PBS rinses, and then cryoprotected and embedded using the same protocol described above. Sagittal sections were cut at 14μm and collected on slides for subsequent staining. β-gal staining was done according to the manufacturer’s recommendations.

### qPCR

To confirm a reduction of Mllt11 transcripts, RNA was extracted from cortices of three controls and cKOs at E18.5 using the RNeasy Micro kit (QIAGEN). RNA was reverse transcribed to cDNA using the SuperScript II Reverse Transcriptase kit (Invitrogen). qPCRs were conducted using the SensiFAST SYBR No-ROX kit (Bioline) using primers previously described (Stanton-Turcotte et al., 2022).

### 5-ethynyl-2’-deoxyuridine *in vivo* Labeling

Dams were injected IP with 30 mg/kg body weight of 5-ethynyl-2’-deoxyuridine (EdU; Invitrogen) 12 hours prior to harvesting at E18.5. Sections were immunostained using the Click-iT kit according to the manufacturer’s protocol (Invitrogen). For co-stains with EdU, the immunohistochemistry (IHC) protocol was adapted such that EdU staining was performed before the addition of the secondary antibody.

### Co-Immunoprecipitation and Western Blot

Co-immunoprecipitation (co-IP) experiments were performed as previously described (Stanton-Turcotte et al., 2022). Briefly, Human Embryonic Kidney (HEK) 293 cells were transfected with either *Myc* tag control or *Myc-Mllt11* constructs, cultured for 24 hours, and lysed for IP using anti-c-Myc agarose resin (Pierce) according to manufacturer’s recommendations. Eluted proteins were resolved on SDS-PAGE gels, transferred on to PVDF membranes as described below, and probed with rabbit anti-NMIIB/Myh10 (1:2000, Bethyl Laboratories) antibodies. For Western blots, lysates were prepared from E18.5 whole brains from 4 control and 5 *Mllt11* cKOs. Samples from the co-IP and were separated on 8% SDS-PAGE gels for 1 h at 120 V and transferred overnight at 20 V on to PVDF membranes (Bio-Rad). Blots were probed with rabbit anti-NMIIB/Myh10 (1:2000, Bethyl Laboratories) and mouse anti-GAPDH (1:1000, ThermoFisher). Secondary antibodies were goat anti-rabbit HRP (1:5000, Invitrogen) and goat anti-mouse HRP (1:5000, Invitrogen). Blots were developed with Clarity Western ECL Substrate (Bio-Rad) and imaged on a ChemiDoc Touch Gel Imaging System (Bio-Rad). Band densitometry was done using Image Lab Software (Bio-Rad).

### Microscopy

Histological images were captured using either a Zeiss AxioObserver fluorescence microscope equipped with an Apotome 2 structured illumination device, 20x and 40x oil immersion objectives, and a Hamamatsu Orca Flash v4.0 digital camera or a Zeiss LSM 710 confocal microscope. β-gal and *Mllt11 in situ* staining was captured using an upright Zeiss PrimoStar compound microscope with an ERc5s color camera. Images were processed using Zen software (Zeiss) and figure montages assembled in Photoshop CS6 (Adobe) and Affinity Photo 2 (Serif).

### Image Sampling and Quantification

For analysis of immunostaining markers and EdU labeling, counting frames were randomly placed in respective layers of interest (RL: 50 x 50μm, EGL: 25 x 25μm and 25 x 50μm, anchoring center base: 25 x 50μm, molecular layer or ML: 75 x 75μm, granular layer or GL: 75 x 75μm, cardinal lobes: 100 x 100μm) using ImageJ (FIJI) (Schindelin et al., 2012). One to eight counting frames were analyzed per histological section, with at least three histological sections of the cerebellum taken from three to five different animals for each strain. Pax6^+^ and EdU^+^ cell counts were performed in entire folia which was partitioned by drawing a straight line between the base of adjacent fissures that make up a single lobe. The “directionality” plugin in FIJI was used to analyze dispersion and orientation of Nestin^+^ Bergmann glia (BG) fibers at anchoring centers. The analysis frame (40 x 40μm) was placed with the upper margin along the midline of the fissure and lateral margin at the base of the fissure along the outer EGL boundary. For directionality analyses, the ‘direction (°)’ value reports the center of the gaussian distribution, and the ‘dispersion (°)’ value reports the standard deviation of the gaussian. The “selection brush tool” in ImageJ was used to outline the entire cerebellum, EGL, and first 100μm of the RL and the area calculated. The “straight” tool in ImageJ was used to measure EGL thickness, GL thickness, ML thickness, fissure depth, and folia height. To measure folia heights, a straight line was drawn from the base of adjacent fissures parallel to the EGL surface. At the midpoint, a perpendicular line was drawn out to the crown of the folia (including the EGL) and a length measurement was taken (Fig. 4K). To measure fissure depths, a straight line was drawn across the crowns of adjacent folia. Along the line (at the fissure point), the “straight” tool was used to measure the depth to the base of the fissure (inner EGL boundary, Fig. 4K). At least 3 axial levels were analyzed per animal. To ensure consistency among samples, cell counts were restricted to images taken from approximately the same axial regions, identified by continuity with the midbrain and ventricular space between the brainstem anteriorly.

### Statistical Analysis

GraphPad Prism V9 software was used to perform all statistical analyses and to produce graphical representations of the data. Statistical differences were determined with Student’s t-tests (two-tailed) with Welch’s correction. In all statistical analyses, a minimum of three control and three cKO animals were used for quantifications using the unbiased and systematic sampling method described previously (Yamada et al., 2015). Bar and line charts were constructed using GraphPad Prism V9 software with results shown as mean ± standard deviation (SD) and each point representing an individual (averaged over three serial images). In all quantification studies, significance level was set at P ≤ .05 (*P ≤ .05, **P ≤ .01, ***P ≤ .001, ****P ≤ .0001).

## RESULTS

### *Mllt11* Expression Pattern in the Embryonic Cerebellum

Previous studies have described *Mllt11* expression in post-mitotic neurons throughout the developing CNS, including the cortex and retina, but its expression in the developing cerebellum remains uncharacterized (Blommers et al., 2023; Stanton-Turcotte et al., 2022; Yamada et al., 2014). To determine if the role of *Mllt11* in neuronal migration generalized to other CNS areas, we began by exploring whether *Mllt11* is expressed during mouse cerebellar development by using the targeted *Mllt11* allele, which includes a *lacZ* gene inserted into the locus. We observed β-gal staining, reflecting *Mllt11* locus expression, in the rhombic lip region from E12.5 to E18.5 (Fig. 2A-E). At E12.5, β-gal staining was observed throughout the dorsal half of the cerebellar primordium (Fig. 2A), exhibiting spatial patterning consistent with the subpial stream of rostrally-migrating Deep Cerebellar Nuclei (DCN) as they migrate into the Nuclear Transitory Zone (NTZ) (Fink et al., 2006). By E14.5, β-gal staining became more widespread across the entire cerebellar primordium, including the developing EGL (Fig. 2B). At this stage, granule cell progenitors (GCPs), which remain in the RL until E13, rapidly emerge from the RL, migrate tangentially to cover the dorsal surface of the primordium, and form the EGL by E15.5 (Capaldo & Iulianella, 2016; Consalez, Goldowitz, Casoni, & Hawkes, 2020; Iulianella et al., 2019). At E16.5, β-gal staining intensity increased and exhibited a similarly widespread distribution throughout the entire cerebellum, including the RL and faintly in the EGL (Fig. 2C). At this stage, GCs continue their tangential migration along the dorsal surface of the cerebellar primordium, with some undergoing radial migration. These radially migrating GCs move inward along Bergmann glial (BG) fibers to populate the cerebellar core in a diffuse manner (Chung, Kim, Jung, Lee, & Jeong, 2010). It is important to note that these can be considered “early” inwardly migrating GCs, as most radial migration occurs after birth. At E18.5, *Mllt11* locus activity was widespread throughout the entire cerebellum, including the RL and EGL (Fig. 2E). At this timepoint, GCs undergo clonal expansion in the EGL, and some postmitotic GCs migrate inward after a switch to radial migration (Chung et al., 2010; Consalez et al., 2020).

**Figure 2.**
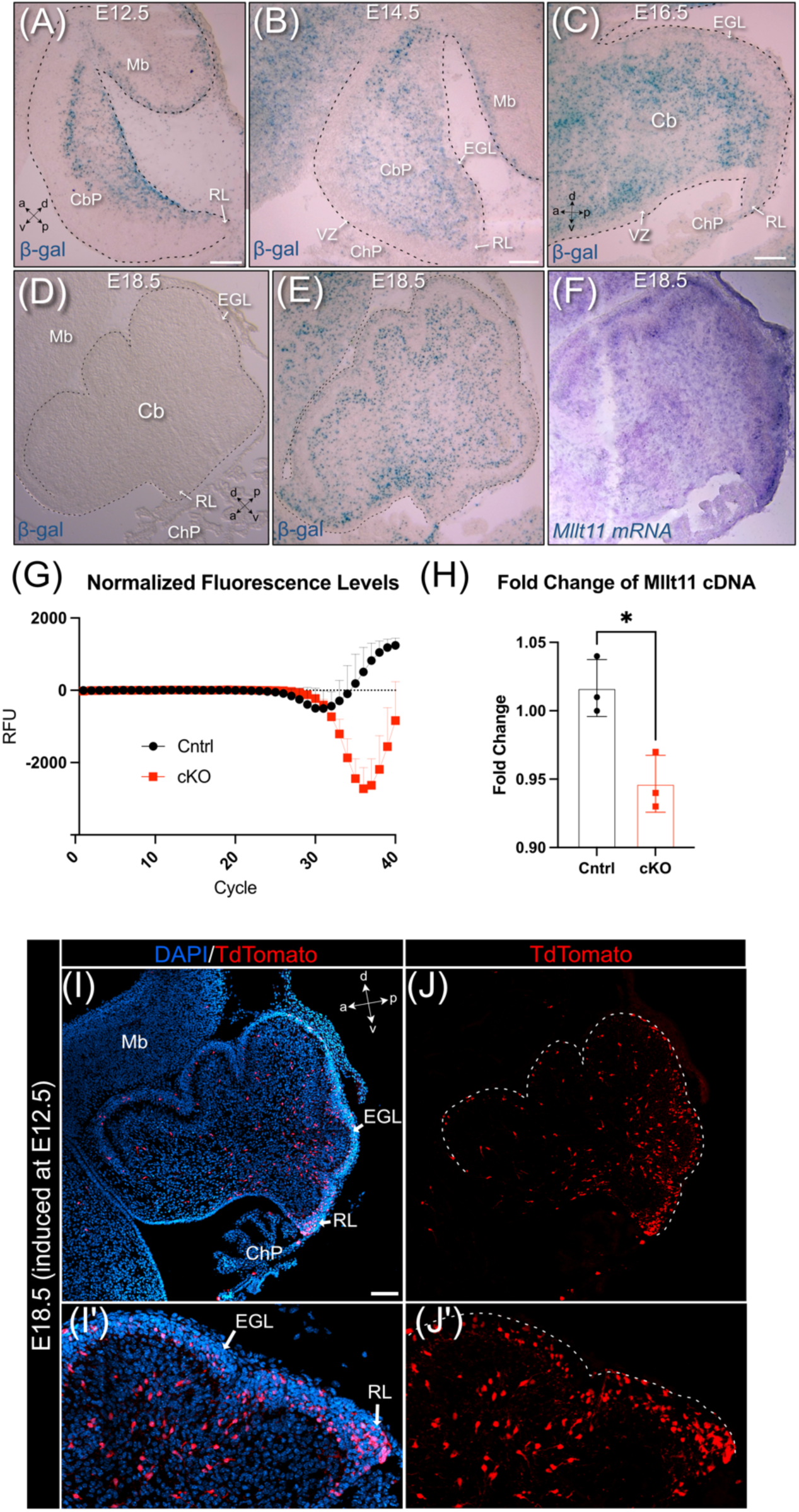
*Mllt11* expression profile and *Cux2^+^* tdTomato-expressing fate mapped cells following tamoxifen induction in the developing cerebellum. (A-F) Sagittal sections taken of the targeted *Mllt11* locus with inserted *LacZ* cDNA (A-C, E) or negative control (D) visualized by β-gal staining, and *Mllt11* mRNA by ISH (F). **(A)** At E12.5, β-gal staining is widespread throughout the dorsal half of the Cbp and faintly in the RL **(B)** At E14.5, β-gal staining was more widespread through the entire primordium including the EGL. **(C)** At E16.5 β-gal staining was intense throughout the core and faintly detected in the RL and EGL. **(D)** Negative control section displaying lack of background expression. **(E)** Medial sections at E18.5 show β-gal staining throughout the core, EGL, and RL. (F) *Mllt11* mRNA expression in the cerebellum at E18.5 reveals a similar expression pattern to the β-gal staining in **(A-C, E)**. **(G)** qPCR fluorescence level quantification of control and *Mllt11* cKO E18.5 cortical tissues normalized to internal control GAPDH **(H)** Significant decrease in fold change of *Mllt11* cDNA transcript levels in cKO relative to controls. **(I-J)** E18.5 parasagittal cerebellum sections from animals induced at E12.5 co-stained with DAPI showed restricted tdTomato^+^ cell fate labelling in the RL, EGL, and some cells in the core. **(I’-J’)** Higher magnification views of the RL region of the corresponding G/H panels. Abbreviations: Cb, Cerebellum; CbP, cerebellar primordium; ChP, choroid plexus; EGL, external granular layer; Mb, midbrain; RL, rhombic lip; VZ, ventricular zone.

To validate the accuracy of β-gal staining in reflecting *Mllt11* gene expression, we conducted ISH on E18.5 cerebella to examine the distribution of *Mllt11* transcripts. The results showed a similarly diffuse distribution of *Mllt11* mRNA throughout the cerebellum, including the RL and EGL (Fig. 2F). Overall, the spatiotemporal expression pattern of *Mllt11* is consistent with the migration of cells in the developing cerebellum, including GCs. We conducted β-gal staining on E18.5 wild type (non-transgenic) cerebellar tissue as a negative control and demonstrate a lack of background staining (Fig 2D), demonstrating the specificity of the targeted *Mllt11* locus activity.

### Conditional Knockout of *Mllt11* in Migrating Granule Cells

The expression pattern of *Mllt11* in the developing RL, EGL, and GL suggested its potential involvement in the migration of GCs (Consalez et al., 2020). To investigate the role of *Mllt11* in the development of migratory GCs, we employed a conditional loss-of-function using a tamoxifen-inducible *Cux2^CreERT2^* knock-in allele, which is restricted to the RL, EGL and most excitatory lineages of the cerebellum, including GCs and unipolar brush cells (UBCs) (Capaldo & Iulianella, 2016). Importantly, our previous work on mapping tamoxifen-inducible Cux2^+^ fated cells demonstrated strong activity in GCPs as well as mature GCs (Capaldo & Iulianella, 2016). Thus, the *Cux2^CreERT2/+^* driver is an ideal genetic strategy to inactivate *Mllt11* in the highly migratory GC population while leaving the VZ derivatives largely unaffected (Capaldo & Iulianella, 2016; Iulianella et al., 2019). The inducible *Cux2^CreERT2^* driver also provides added temporal control over *Mllt11* excision, allowing for the analysis of each GC-specific developmental process (Fig. 2G-H). To visualize Cre expressing cells in cKO and control mice, the *Ai9 TdTomato* reporter transgene was utilized, which exhibited tdTomato^+^ fluorescence upon Cre-mediate excision. Analyses were performed at E18.5 when the cerebellar surface has undergone morphological changes, including the formation of the four principal fissures (Preculminate (Pc), Primary (Pr), Secondary (Sec), and Posterolateral (Pl) that separate the five cardinal lobes (Anterobasal (Ab), Anterodorsal (Ad), central, posterior, and inferior; Fig. 1F) (Altman & Bayer, 1985a, 1985b; Miale & Sidman, 1961). To capture GCPs as they leave the RL, form the EGL, and begin to migrate into the core, tamoxifen was administered at E12.5. Cre-mediated excision of *Mllt11* at this time point allowed for investigation of its function in tangential migration of GCPs as they form the EGL as well as radial migration of early, inwardly migrating GCs (Fig. 1B, D).

To corroborate the reduction in *Mllt11* expression in *Cux2*-expressing tissues, qPCR was performed on cortical lysates revealing fold change of *Mllt11* cDNA transcript levels were significantly decreased in cKOs relative to controls (Fig. 2G-H). Cell-specific Cre activity in the developing cerebellum was confirmed with tdTomato fluorescence in the RL, EGL, and some cells within the core of E18.5 cerebella (Fig. 2G, H, G’, H’), consistent with previous work using this genetic strategy (Capaldo & Iulianella, 2016).

To confirm the identity of tdTomato^+^ cells, we stained for cell-type specific markers: Pax6 for GC, Calbindin for Purkinje cells, and Pax2 for inhibitory interneurons (Fig. 3C). Nearly all tdTomato-labeled cells were Pax6^+^ GCs (Fig. 3A-B, D-E) in both the RL (Fig. 3B’, D’, E’) and core (Fig. 3B’’, D’’, E’’) regions of the cerebellum. No tdTomato^+^ cells expressed Pax2 (Fig. 3F-H) and few exhibited cytoplasmic Calbindin staining (Fig. 3I-K). At E18.5, Purkinje cells localize to a diffuse region underneath the EGL, called the Purkinje Plate (PP), before progressively settling into a monolayer postnatally. TdTomato^+^ cells found within the PP likely correspond to GCs that migrated over, under, or between Purkinje cells, as there was minimal overlap between TdTomato^+^ and Calbindin^+^ staining in most cells (Fig. 3I-K).

**Figure 3.**
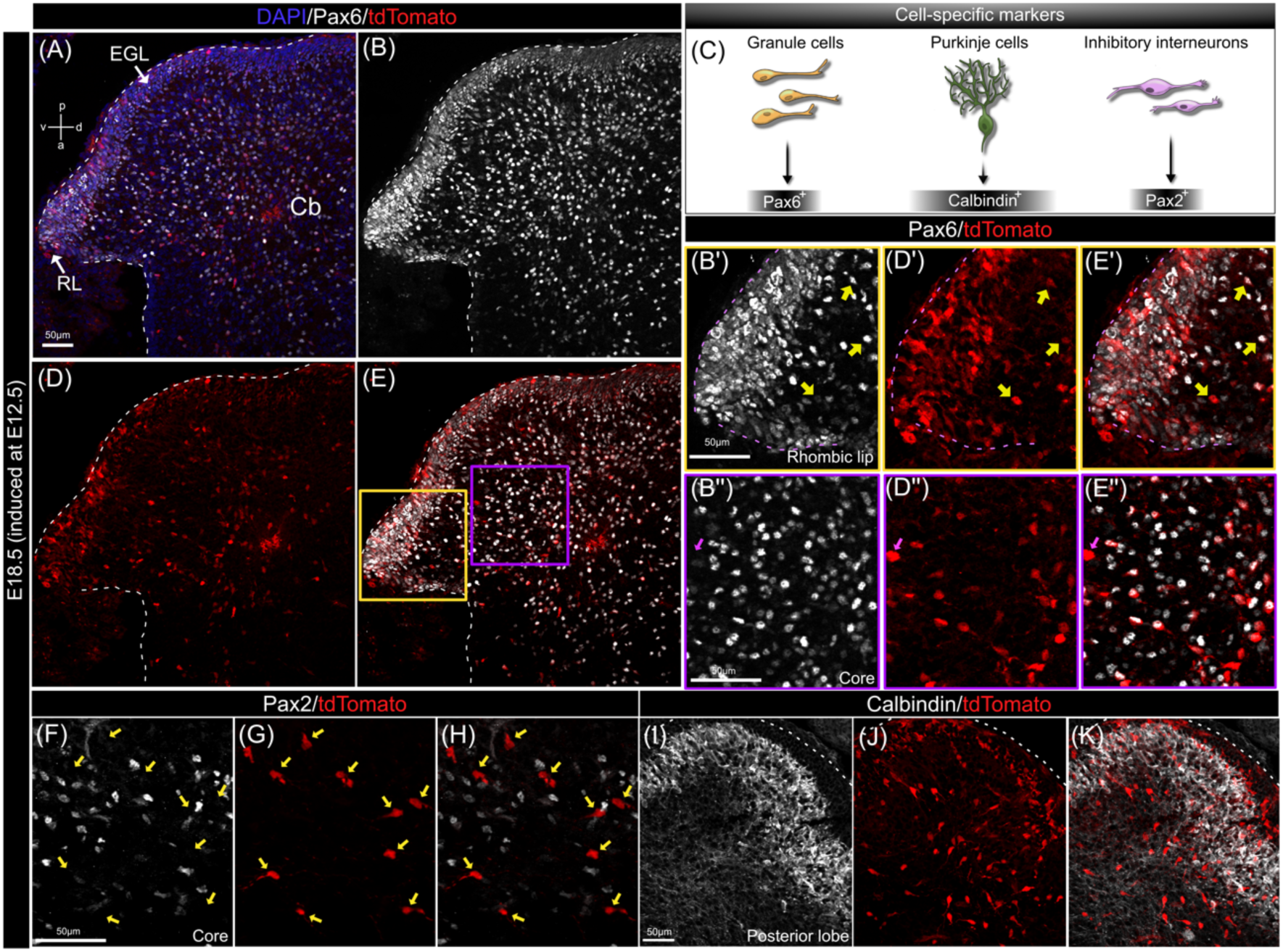
*Cux2^CreERT2^*-driven recombination in granule cells. **(A-B, D-E)** Parasagittal section of an E18.5 cerebellum induced at E12.5 exhibited tdTomato^+^-labeled cells co-expressed with the GC marker, Pax6. Yellow and purple insets in **(E)** correspond to color-coordinated borders of panels **(B’, D’, E’, B’’, D’’, E’’)**. **(B’, D’, E’)** Almost all tdTomato^+^ cells in the RL and adjacent EGL were Pax6^+^. **(B’’, D’’, E’’)** Most tdTomato^+^ cells in the core expressed Pax6 except a few which were Pax6^-^ (fuchsia arrow). **(C)** Schematic of cerebellar cell-type specific transcription factors markers used in the study. **(F-H)** TdTomato-labeled cells in the core of the cerebellum did not express the inhibitory interneuron marker Pax2 (yellow arrows). **(I-K)** Sample image from the posterior lobe revealed tdTomato^+^ cells did not express the Calbindin. Most tdTomato^+^ cells seen in the Calbindin+ PP were inwardly migrating GCs passing thorough the PP. Scale bar: 50μm for **(A-K)**. Abbreviations: a, anterior; d, dorsal; Cb, cerebellum; EGL, external granular layer; p, posterior; RL, rhombic lip; PP, Purkinje plate; v, ventral.

### *Mllt11* Mutant Cerebella Exhibit Decreased Area, Shallower Principal Fissures, Smaller Folia, and Enlarged Rhombic Lips

Gross evaluation of *Mllt11* cKOs at E18.5 revealed smaller cerebella with significantly reduced foliation compared with controls (Fig. 4A-B). Total cerebellar area was significantly decreased in *Mllt11* mutants (Fig. 4A-B, C; *P* = 0.002, N = 5), accompanied by a reduction in EGL areas (Fig. 4A-B, C; *P* = 0.002, N = 5). When expressed as a ratio (EGL to total cerebellar area), there was no difference between genotypes (Fig. 4C; *P* = 0.653, N = 5), indicating a proportional reduction in EGL size relative to the decrease in total cerebellar area observed in *Mllt11* cKOs.

**Figure 4.**
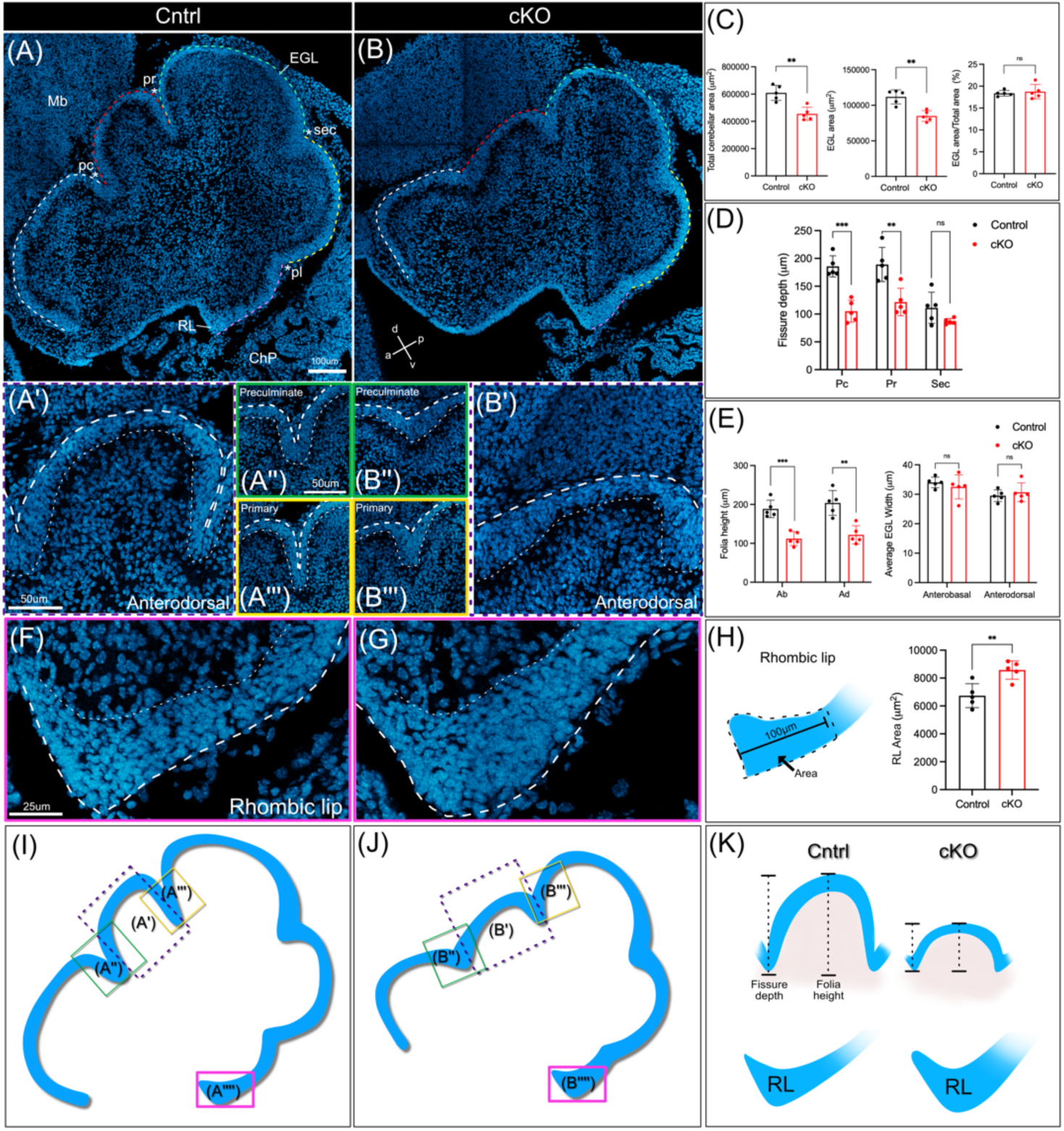
*Mllt11* loss resulted in smaller cerebella with shallower principal fissures, smaller folia, and enlarged Rhombic Lip. **(A, B)** Parasagittal sections of E18.5 cerebella stained with DAPI revealed controls **(A)** with larger cerebellar and EGL areas compared to *Mllt11* cKO mutants **(B)**. **(A, B)** At E18.5 the four principal fissures and five cardinal lobes are present in both controls **(A)** and mutants **(B)**: Ab (white outline), Ad (red outline), central (green outline), posterior (yellow outline), and inferior (pink outline). **(C)** Bar chart comparisons of total cerebellar area, EGL area, and respective ratios for controls versus *Mllt11* cKOs. **(A’)** Ad lobes were taller in controls compared to cKOs **(B’)**. Pc **(A’’, B’’)** and Pr **(A’’’, B’’’)** fissure depths were also deeper in controls **(A’’, A’’’)** compared to cKO mutants **(B’’, B’’’)**. Colored outlines in **(A’-A’’’, B’-B’’’, F, G)** correspond with insets in underlying panels. **(D)** Bar chart comparisons of three principal fissure (Pc, Pr, and Sec) depths **(E)** Bar chart comparisons of Dorsal folia (Ab and Ad) heights, and average EGL widths in controls vs. cKOs. **(F)** Control RLs were smaller than **(G)** cKOs. **(H)** Schematic of how RL areas were calculated (first 100μm) with a bar chart comparison of RL areas of controls vs. cKOs. **(I, J)** Schematic EGL tracings in control **(I)** and cKO **(J)** cerebella with inset boxes color-coordinated to outline corresponding panels above. **(K)** Schematic summary of control versus *Mllt11* cKO comparing folia height, fissure depth, and RL size. Welch’s t-test, **(A, B, F, G)** N = 5. Data presented as mean ± SD. N.s., not significant. **P ≤ .01; ***P ≤ .001. Scale bar: 100μm for **(A, B)**, 50μm for **(A’-A’’’, B’-B’’’)**, and 25μm for **(F, G)**. Abbreviations: a, anterior; Ab, anterobasal; Ad, anterodorsal; ChP, choroid plexus; d, dorsal; EGL, external granular layer; Mb, midbrain; p, posterior; pc, preculminate; pl, posterolateral; pr, primary; RL, rhombic lip; sec, secondary; v, ventral.

*Mllt11* cKOs exhibited shallower depths of both the Preculminate (green boxes in Fig. 4A’’, vs B’’, D; *P* = 0.0002, N = 5) and Primary (yellow boxes in Fig. 4A’’’, vs B’’’, D; *P* = 0.006, N = 5) fissures compared with controls. Although statistical analysis of the Secondary fissure did not reach significance (Fig. 4A, B, D; *P* = 0.125, N = 5), *Mllt11* mutants still displayed a noticeable reduction in its depth. In conjunction with the shorter fissures, *Mllt11* cKOs exhibited shorter anterobasal (Ab, Fig. 4A-B, E; *P* = 0.0004, N = 5) and anterodorsal (Ad, Fig. 4A’ vs B’, E; *P* = 0.002, N =5) cardinal lobes. The average width of the EGL did not differ markedly between *Mllt11* cKOs and controls at the crown of either lobe, while folia height, which included the EGL in measurements, were significantly shorter in cKOs (Fig. 4E, K for diagram of height measurement). During foliation, anchoring centres at the base of each fissure remain relatively fixed while folia grow outward. All four principal fissures formed in *Mllt11* cKOs, but the Preculminate and Primary were significantly shallower compared to controls (Fig. 4D), indicating potential disruptions in folia lengthening once anchoring centres had been established. Folia lengthening is a complex process driven by self-sustaining proliferation of GCPs and their directed radial migration along BG fibers, suggesting that *Mllt11* loss may impact radial migration, the pre-existing folia cytoarchitecture, or both (Sudarov & Joyner, 2007).

Interestingly*, Mllt11* cKO cerebella also exhibited enlarged RLs relative to controls (Fig. 4F-G, H, K; *P* = 0.006, N = 5). The RL region is densely packed with GCPs that have yet to migrate tangentially to form the EGL, implying that *Mllt11* may play a role in the tangential migration of GCs from the RL and its inactivation results in a migration deficit and retention of GCPs unable to migrate. Taken together with the role of Mllt11 in neuronal migration in the neocortex and retina, the altered folia morphology and thickening of the RL imply a migration deficit (Blommers et al., 2023; Stanton-Turcotte et al., 2022).

### *Mllt11* Loss Leads to Impaired Granule Cell Migration

GCs rely on a combination of intrinsic and extrinsic programs to regulate their migration, in the form of morphological changes mediated by the cytoskeleton and contact with underlying glial scaffolding to initiate glial-guided migration, respectively. Because GCs undergo both radial and tangential migration along their path from the RL to the EGL and cerebellar core (Fig. 1B, D), it is possible that a migration deficit could be the result of impairments to either migratory mode or both concurrently. As tangential migration temporally precedes the initiation of radial migration and positions GCs in the EGL before they invade the core via glial fibers, it is possible that an initial tangential migratory deficit could contribute to the altered radial distribution of GCs, causing morphological deficits in the folia. We therefore decided to investigate the potential tangential migration defect exhibited by *Mllt11* cKOs by pulsing pregnant dams with EdU late on E17 and analyzing the localization of EdU labelled cells 12 hours later. As EdU is incorporated into DNA of cells and their progeny, dosage at this time point allowed for labelling of the final wave of tangentially migrating GCPs from the RL (Fig. 6G, H, K, L) (Rahimi-Balaei et al., 2018).

**Figure 5.**
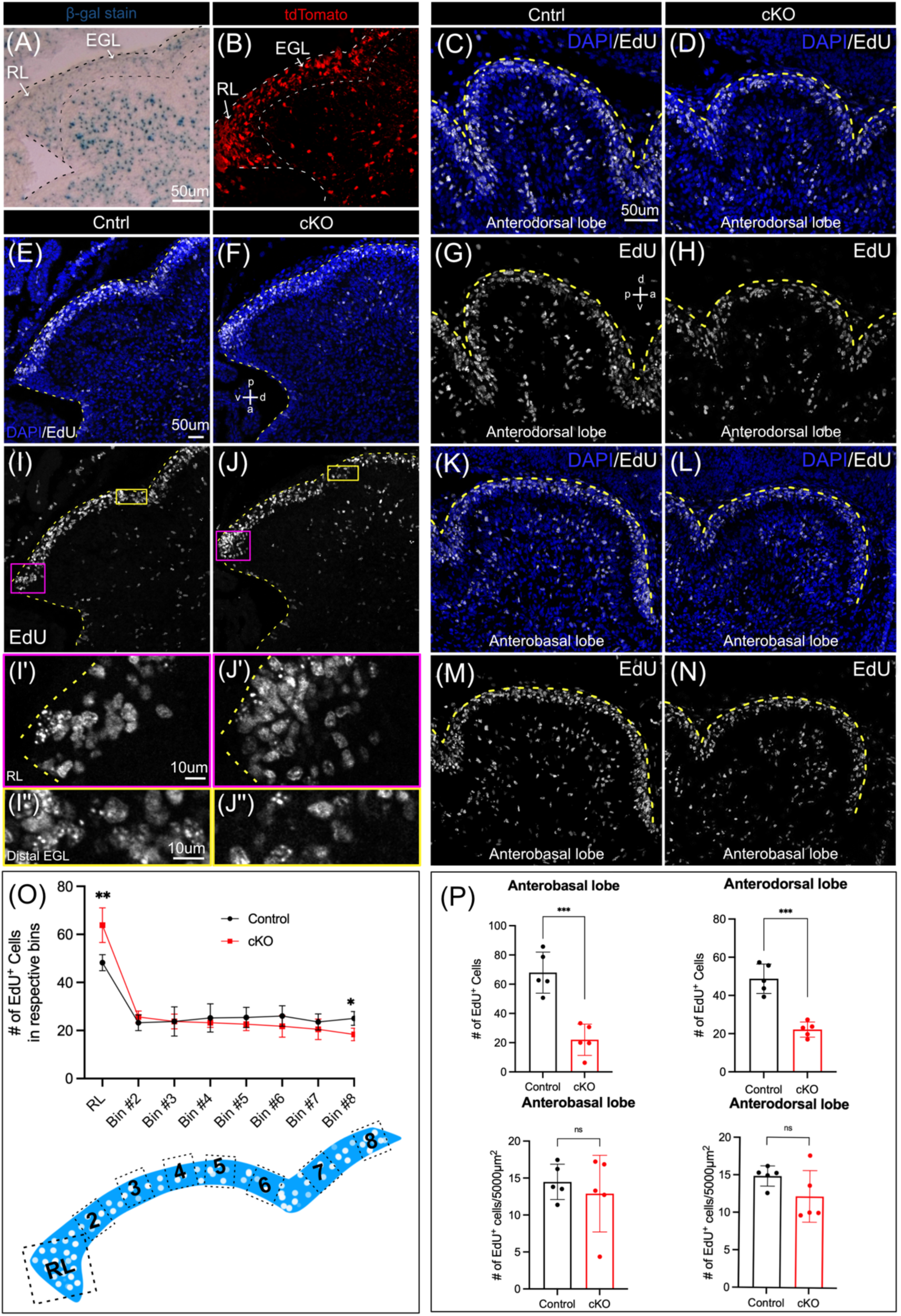
*Mllt11* cKO mutants exhibit reduced EdU^+^ granule cell precursor migration from the Rhombic Lip. **(A)** β-gal staining in the RL region revealing *Mllt11* locus activity in the RL, EGL, and cerebellar anlage core at E18.5, which overlapped where *Cux2^CreERT^*^2^-mediated activation of tdTomato^+^ reporter gene fluorescence **(B)** induced at E12.5. **(C)** Control and **(D)** *Mllt11* cKO Anterodorsal (Ad) lobes co-stained with DAPI. **(E-F)** EdU^+^ cell labeling in the RL region including the inferior and posterior cardinal lobes, co-stained with DAPI. **(G-H)** EdU^+^ nuclei labeling reveals a larger cohort of cells populating the Ad lobe in **(G)** controls verses **(F)** *Mllt11* cKOs. **(I-J)** EdU^+^ nuclei in the RL and EGL of **(I)** control and **(J)** *Mllt11* cKOs. Color-coded insets corresponding to panels I’-J”. **(I’)** Fewer EdU^+^ nuclei in the RL of controls vs. cKOs **(J’)**. **(I’’)** Distal EGL of controls exhibited a greater number of EdU^+^ nuclei versus cKOs **(J’’)**. **(K)** Control and **(L)** cKO Anterobasal (Ab) lobes co-stained with DAPI. **(M-N)** EdU^+^ nuclei labeling shows a larger cohort of cells populating the Ab lobe in **(M)** controls vs. **(N)** cKOs. **(O)** Line chart comparison of EdU^+^ cells in bins placed in the RL (50 x 50mm) and EGL region (25 x 25mm) moving distally in controls (black) vs. cKO (red). Below is a schematic representation of the bin areas corresponding with the cell counts along the x axis. **(P)** Graphical comparison of EdU^+^ nuclei in the Ab and Ad lobes of controls (black) verses cKOs (red) with counts normalized per 5000mm^2^ area. Welch’s t-test, N = 5. Data presented as mean ± SD. n.s., not significant; ** = p ≤ .01; *** = p ≤ 0.001. Scale bar: 50μm (A-F), 10μm for (I’-J’’). Abbreviations: a, anterior; d, dorsal; EGL, external granular layer; p, posterior; RL, rhombic lip; v, ventral.

**Figure 6.**
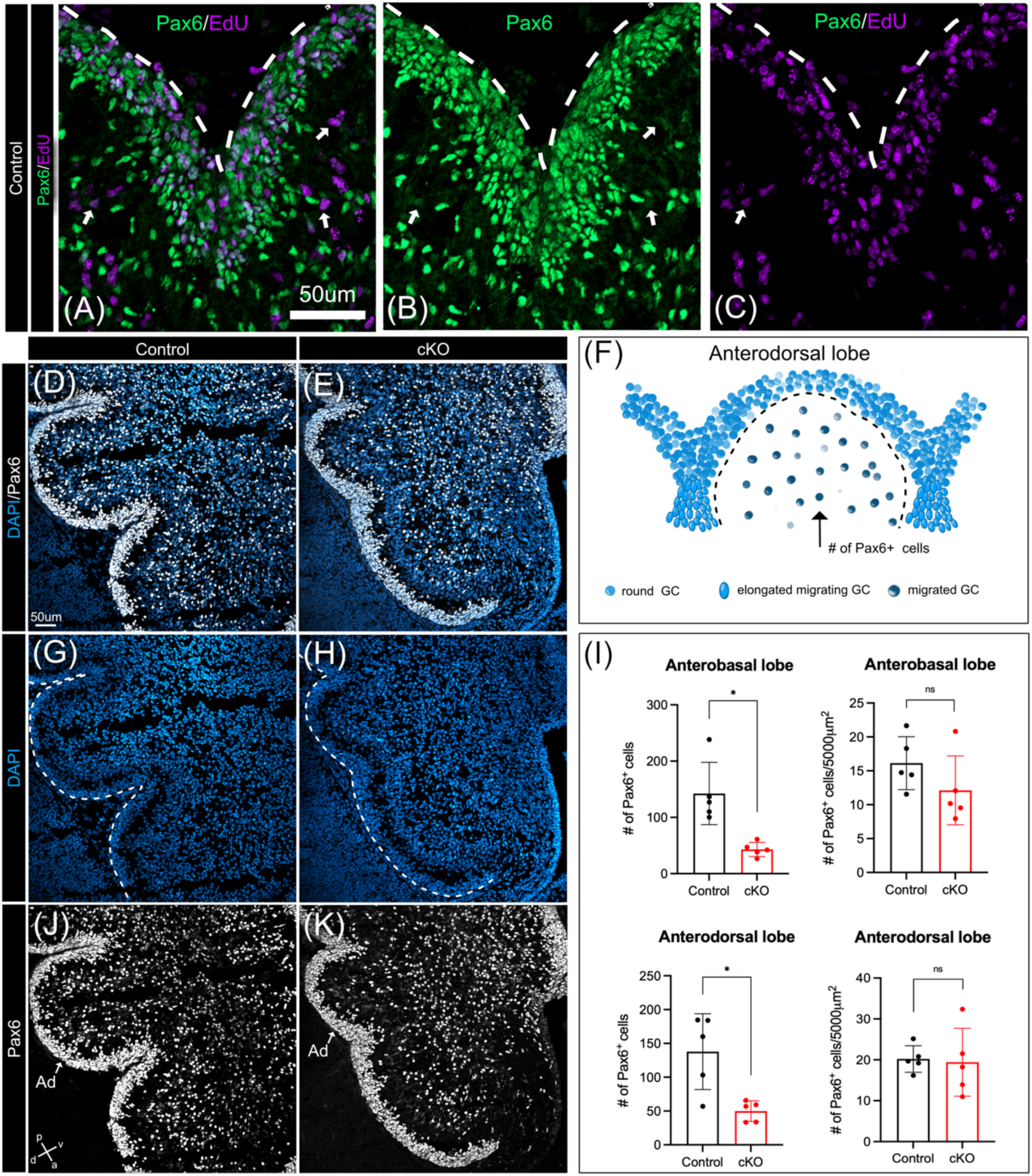
*Mllt11* cKO mutants exhibit reduced invasion of granule cells at folia anchor points. **(A-C)** EdU and Pax6 immunohistochemistry in a control section, exhibiting EdU^+^/Pax6^+^ co-stained nuclei, identified by white arrows, indicating inwardly migrating GCs. **(D-E, G-H, J-K)** Medial sagittal sections of E18.5 cerebella stained with Pax6 and DAPI. Images were taken to capture the dorsal cardinal lobes and cerebellar core. **(D, G, J)** Sections from control cerebella show large folia and deep fissures in comparison to *Mllt11* cKO **(E, H, K)** cerebella which show shorter folia and shallower fissures. **(I)** Bar charts displaying comparisons of control (black) versus cKO (red) Pax6^+^ counts **(F)** Schematic of the Anterodorsal lobe to illustrate how the Pax6^+^ counts in **(I)** were performed. Welch’s t-test, **(I)** N = 5. Data presented as mean ± SD. n.s., not significant; ** = p ≤ .01; *** = p ≤ 0.001. Scale bar: 50μm. Abbreviations as in Fig. 5.

Since *Mllt11* was expressed in the RL and adjacent EGL at E18.5 (Fig. 5A), and the excision event was restricted to this region (Fig. 5B), this genetic strategy and birth-dating experiment provided an ideal approach to investigate potential tangential migration defect of GCPs emanating from the RL. As GCPs are born in the RL and migrate rostrally over the dorsal surface of the developing cerebellum, we analyzed the RL and seven sequential regions of the EGL to follow EdU^+^ cells along their migratory path (Fig. 5O). In the RL, controls (Fig. 5I’) displayed fewer EdU^+^ cells relative to cKOs (Fig. 5J’), which was more densely packed with labelled cells (Fig. 5O; *P* = 0.005, N = 5). Moving along sequential bins there was a progressive, non-significant trend toward fewer EdU^+^ cells in cKOs (Fig. 5I, J). Analysis of the distal-most EGL, bin (#8) revealed a significant reduction in EdU^+^ cells in cKOs (Fig. 5J’’) relative to controls (Fig. 5I’’, P; *P* = 0.005, N = 5). This altered distribution of GCPs across their tangential migratory path in *Mllt11* cKOs compared with controls suggest impaired tangential migration.

It is important to consider that GCP proliferation in the EGL temporally coincides with the developmental period during which they migrate tangentially. Therefore, it is possible that *Mllt11* loss led to a reduction in EdU^+^ cells in the distal EGL either by affecting the migration or proliferation of GCPs during or after their migration, or both. To address this, we took advantage of the temporal overlap between the final wave of tangential GCP migration from the RL at E17.5 and radial migration of GCs into the cerebellar core by labeling this late cohort of GCs with an EdU pulse at E17. This will capture cells within the RL, EGL, and core of control (Fig. 5C, G, K, M) and *Mllt11* cKO cerebella (Fig. 5D, H, L, N) (Chung et al., 2010). *Mllt11* cKOs exhibited a clear reduction in the number of EdU^+^ cells in both the anterodorsal (Fig. 5H, P; *P* = 0.0005, N = 5) and anterobasal (Fig. 5N, P; *P* = 0.0005, N = 5) lobes compared with controls (Fig. 5G, M). Co-staining with GC marker Pax6 revealed that some EdU^+^ cells populating dorsal folia expressed low levels of Pax6, suggesting they were postmitotic GCs that had migrated inward (Fig. 6A-C, white arrows, F) (Engelkamp, Rashbass, Seawright, & van Heyningen, 1999; Englund et al., 2006).

Pax6 expression is maintained in GC precursors throughout their maturation process, from the RL at birth to their final position in the IGL (Stoykova & Gruss, 1994). At E18.5, Pax6 robustly labeled GCs and their precursors throughout the developing cerebellum, including the EGL and core (Fig. 6D-E, J-K). To further investigate a role for Mllt11 in radial invasion into the cerebellar core, we quantified GCs that had successfully migrated inward from the EGL and localized within the folia (Fig. 6F). We found a significant reduction in the number of Pax6^+^ cells within both the anterobasal (Fig. 6E, H, K, I; *P* = 0.01, N = 5) and anterodorsal (Fig. 6E, H, K, I; *P* = 0.02, N = 5) lobes in *Mllt11* cKOs (Fig. 6D, G, J, I).

Many EdU^+^ cells were Pax6^-^, resulting from the labeling of a wide variety of cerebellar cell types with the E17 pulse, including of Purkinje cells and inhibitory interneurons derived from the VZ (Fig. 6A-C). Normalizing EdU and Pax6 counts to folia area masked the difference in the anterodorsal (Fig. 5P; *P* = 0.167, N = 5, 6I; *P* = 0.85 N = 5) and anterobasal (Fig. 5P; *P* = 0.558, N = 5, 6I; *P* = 0.203, N = 5) lobes between genotypes. Taken together, these finding suggest that the loss of *Mllt11* likely affected radial migration of GCs, resulting in underdeveloped folia with smaller areas (Fig. 5G-H, M-N).

### *Mllt11* Loss Disrupted Bergmann Glia Fibers and Reduced Pax6^+^ cells at Anchoring Centres

Since both tangential and radially migrating cells exhibited altered distributions across their anticipated trajectory in *Mllt11* mutants, it is difficult to parse whether the migratory deficits exhibited were due to cell-intrinsic mechanisms required for movement, or cell-extrinsic defects due to abnormalities in the underlying cytoarchitecture that provides a conduit for cells to travel along. As the cerebellar surface undergoes foliation, BG fibers surrounding the base of each fissure undergo a shift in orientation, transitioning from parallel to radiating inward towards the anchoring centre, serving as tracks for inwardly migrating GCs (Fig. 7E). We visualized BG by immunostaining for Nestin, a type IV intermediate filament protein expressed by neural stem cells and BG (Li et al., 2013). Nestin^+^ fibers exhibited an atypical fanning pattern at anchoring centres in *Mllt11* cKOs relative to controls (Fig. 7A-D). In the Preculminate fissure, *Mllt11* cKOs exhibited parallel-oriented BG fibers lacking proper radiating patterns towards the base of the fissure (Fig. 7B, D). In contrast, control cerebella displayed the typical fanning pattern with fibers radiating towards the base (Fig. 7A, C). Quantitative analysis of BG fiber directionality at the Preculminate fissure revealed a reduction in fiber dispersion (Fig. 7F; *P* = 0.004, N = 4) and average angle (direction; Fig. 7F; *P* = 0.0009 N = 4) in *Mllt11* cKOs relative to controls. Analysis at the Primary fissure showed a non-significant reduction in dispersion (Fig. 7F; *P* = 0.466, N = 4) and average angle (direction; Fig. 7F; *P* = 0.055, N = 4). Examination of inward migration at anchoring centres revealed reduced Pax6^+^ cell counts just inside the Preculminate (Fig. 7G-J; *P* = 0.03, N = 5) and Primary (Fig. 7G-J; *P* = 0.003, N = 5) fissures in *Mllt11* cKOs compared with controls (Fig. 7H-J). Taken together, this suggests *Mllt11* loss led to altered orientation of BG fibers at anchoring centers.

**Figure 7:**
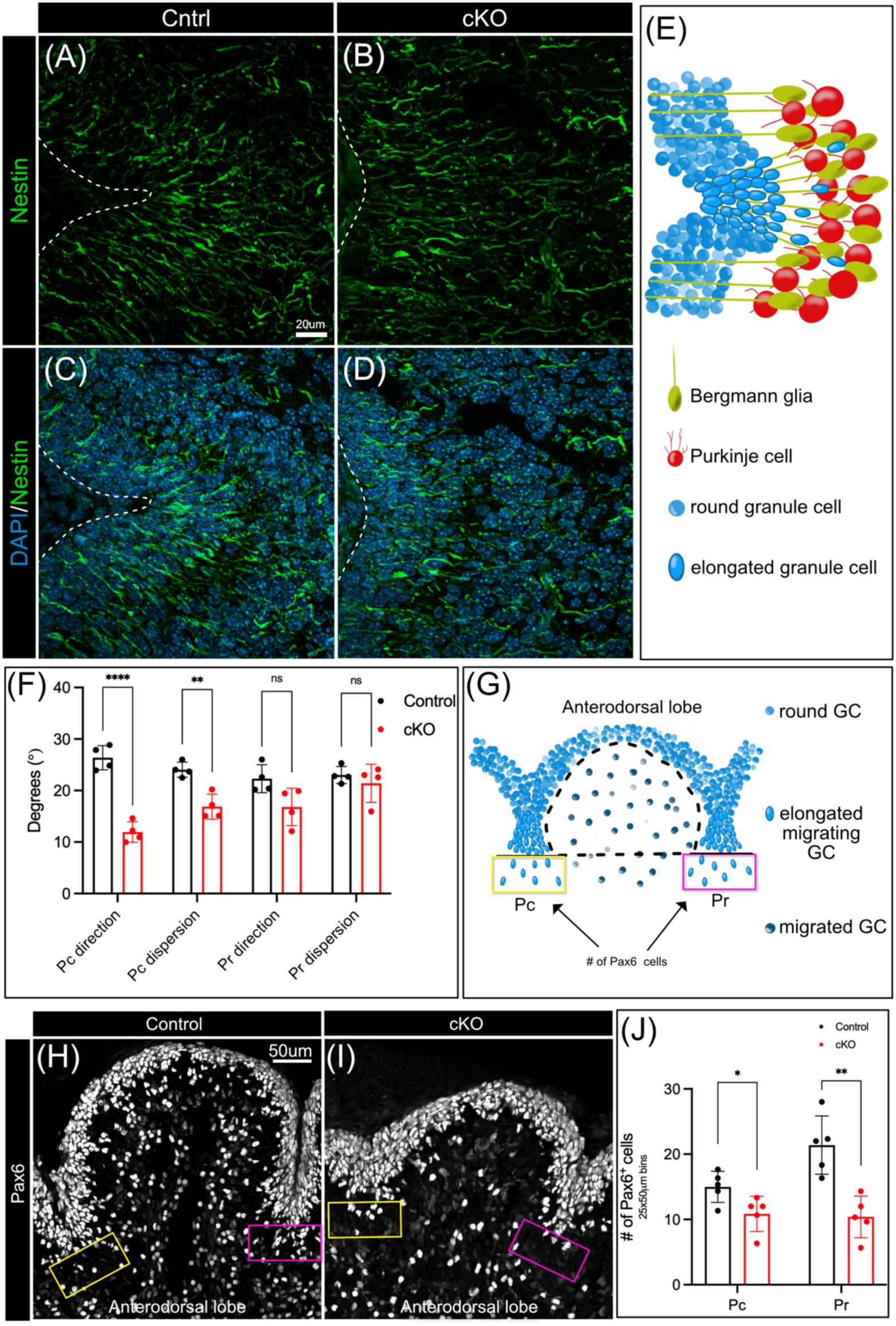
*Mllt11* loss disrupted Bergmann glia fibers at anchoring centers. **(A-D)** Sagittal sections of E18.5 control **(A, C)** and *Mllt11* cKO **(B, D)** cerebella at the Pc fissure stained with Nestin to visualize BG fibers. **(A)** Controls exhibited a typical fanning pattern of BG fibers while cKOs **(B)** had atypical fibers running in parallel. **(E)** Schematic depicting an *anchoring centre* which forms at the base of each fissure and coincides with increased GC proliferation, EGL thickening, invagination of the underlying Purkinje cell layer, and BG fibers radiating towards the base of the fissure in a fanning pattern. **(F)** Bar graph comparisons of average direction (°) and dispersion (°) angle of fibers in a 40 x 40μm bin placed at the base of each fissure with the upper margin of the bin aligning with the midline of the fissure. Controls (black) exhibited typical fanning pattern of BG fibers at anchoring centres with a greater average angle (direction) and degree of dispersion at the Pc fissure. *Mllt11* cKOs (red) exhibited smaller average fiber angles (direction) and degrees of dispersion at the Pc but not the Pr fissure. **(G)** Schematic of the Anterodorsal lobe to illustrate how Pax6^+^ cells were counted. **(H-J)** High magnification views of Pax6 immunohistochemistry the Anterodorsal lobes in controls **(H)** vs. cKOs **(I)**. At each anchoring centre below the EGL Pax6^+^ cells were counted (yellow and pink boxes). **(J)** Bar chart comparison of control (black) vs. cKO (red) Pax6^+^ cell counts in 25 x 25mm bins placed at the base of the Pc and Pr fissures just inside the EGL. Welch’s t-test, (A, B) N = 4. Data presented as mean ± SD. n.s., not significant; ** = p ≤ .01; **** = p ≤ .0001. Scale bar: 20μm for (A-D). Abbreviations: Pc, preculminate; Pr, primary.

### *Mllt11* Loss Left the Purkinje Cell and Inhibitory Interneuron Populations Largely Unaffected

In addition to GCs and BG, Purkinje cells are crucial for anchoring the developing architecture of the cerebellum. While the genetic strategy employed for the current study restricted *Mllt11* loss to predominantly GCs, it was important to investigate any possible effects on other cerebellar populations, namely Purkinje cells and inhibitory interneurons. At late embryonic stages Purkinje cells arrange themselves in a diffuse multilayer Purkinje plate (PP) below the EGL, which undergoes inward folding, corresponding to the thickening and invagination of the overlying EGL, likely due to the accumulation of migrating GCs (Sudarov & Joyner, 2007). Given the disrupted inward migration of GCs and fissure development in *Mllt11* cKOs, we investigated the impact of *Mllt11* loss on the PP. We used Calbindin D-28K immunostaining to visualize Purkinje cells and PP invaginations (Fig. 8A-D) (Barski et al., 2003), and observed that cKOs exhibited a disrupted organization of Purkinje cells in the PP, instead of a columnar cluster typically associated with the wild type controls cerebellum (Fig. 8D’ vs. C’). Although Purkinje cells did not accumulate in subcortical regions, and the PP still formed in cKOs, but the degree of infolding at dorsal fissures was reduced in the mutants (Fig. 8C’-D’).

**Figure 8:**
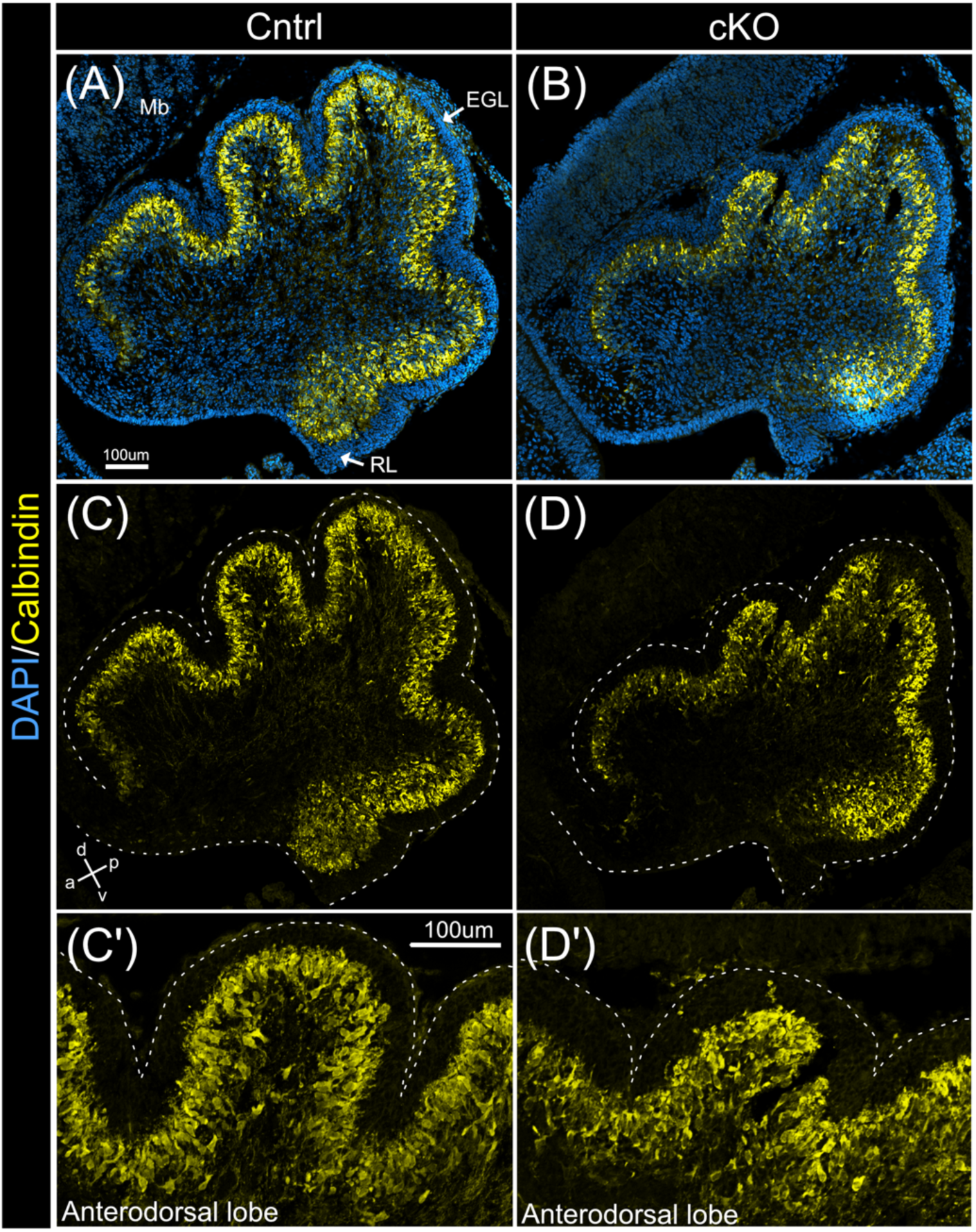
*Mllt11* loss altered the infolding of the Purkinje plate. **(A-D)** Medial sagittal sections of E18.5 cerebella stained with Calbindin to label the PP beneath the EGL which formed in both controls **(A, C)** and *Mllt11* cKOs **(B, D)**. **(A, C)** Controls exhibited a highly invaginated PP at dorsal fissures, while cKOs **(B, D)** were less invaginated. **(C’-D’)** High magnification view of the Ad lobe revealing reduced infolding of the PP in cKOs **(D’)** compared with controls **(C’)**. Scale: 100μm in **(A-D, C’-D’)**. Abbreviations as in Fig.4; PP, Purkinje plate.

The cerebellum also contains inhibitory interneurons, which are generated from the VZ around E13 and subsequently delaminate and migrate into the cerebellar parenchyma (Leto et al., 2016). By late embryonic stages, interneurons are distributed throughout the core and developing folia (Grimaldi, Parras, Guillemot, Rossi, & Wassef, 2009; Weisheit et al., 2006). At E18.5, the inhibitory interneuron marker Pax2 showed reduced distribution in the distal tips of the developing folia of *Mllt11* cKOs in (Fig, 9B, D) relative to controls (Fig, 9A, C) (Weisheit et al., 2006). However, cell counts in each cardinal lobe (Fig. 9E) revealed no significant differences in Pax2^+^ interneurons, except in the anterodorsal lobe, where there were more Pax2^+^ cells in cKOs (Fig. 9C, D, F; *P* = 0.039, N = 4). No significant differences were detected in the numbers of inhibitory interneurons in the remaining cardinal lobes (Fig. 9F). Altogether, the pattern of Pax2 staining reflected the aberrant formation of folia in the *Mllt11* mutant cerebellum, but numbers remained largely unchanged, demonstrating the specificity of the *Cux2^CreERT2^* to the RL and excitatory lineages of the cerebellum.

**Figure 9:**
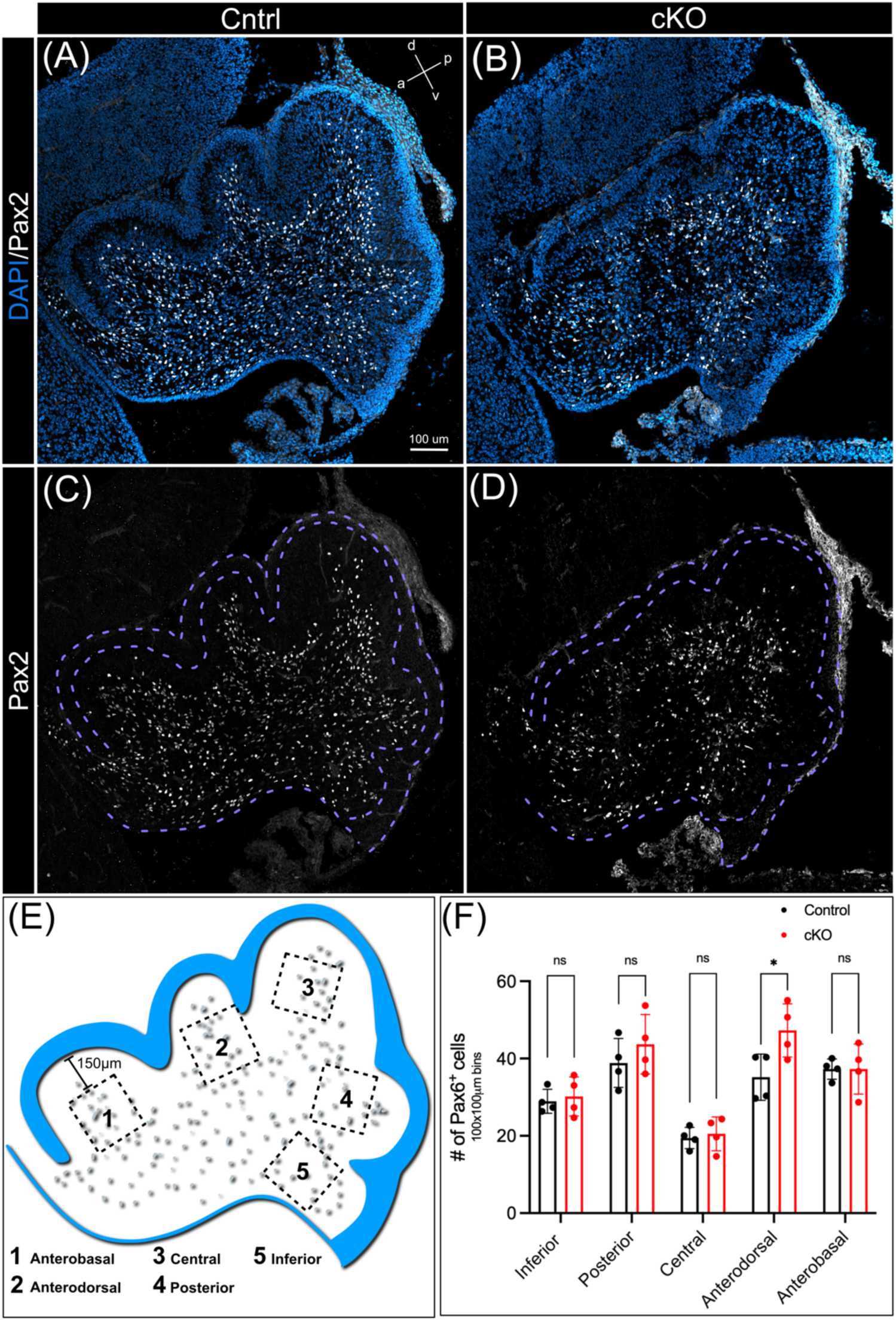
Pax2^+^ interneuron population was largely unaffected by *Mllt11* loss. **(A-D)** E18.5 medial sagittal cerebellar sections with Pax2 staining for inhibitory interneurons. **(A, C)** Control inhibitory INs populated deeper into the developing dorsal lobes compared to **(B, D)** *Mllt11* cKOs. **(E)** Schematic depicting how Pax2^+^ cell counts were performed in each cardinal lobe (150μm from the inner EGL surface). **(F)** Bar chart comparison of Pax2^+^ cells in each identified cardinal lobe in 100 x 100βm bins in control vs. cKOs. No difference was found across lobes except in the Ad, likely reflecting the smaller folia and accumulation of Pax2^+^ cells in the core of cKO cerebella. Welch’s t-test, **(A-D)** N = 4. Data presented as mean ± SD. n.s., not significant; * = p ≤ .05. Scale: 100μm in (A-D).

### *Mllt11* Loss Led to Increased NMIIB Levels

Given the consistency in the apparent migratory phenotype shown here, as well as in the cortex and retina we reported in previous studies (Blommers et al., 2023; Stanton-Turcotte et al., 2022), we investigated whether any of the Mllt11-interacting proteins previously identified in a proteomic screen of fetal brain lysates might be involved in controlling cellular migration. Specifically, in a previously study we identified various tubulins, actin, and non-muscle myosins as possible Mllt11-interacting proteins in the fetal brain (Stanton-Turcotte et al., 2022). In that study we confirmed interaction with tubulins but not myosins. One potential Mllt11-interaction target we decided to focus on was non-muscle myosin IIB (NMIIB/Myh10) because it has been previously implicated in the regulation of cell migration (Ma et al., 2007; Ma et al., 2004).

We first investigated whether IP of myc-Mllt11-transfected HEK293 cells could bring down NMIIB and confirmed that Mllt11 can interact with NMIIB (Fig. 10A). Next, we examined whether *Mllt11* loss impacted NMIIB levels in the brain. For this assay we used E18.5 whole brain lysates from *Cux2^ireCre^; Mllt11* cKOs described in (Stanton-Turcotte et al., 2022) to ensure a robust *Mllt11* knockout in Cux2-expressing cells throughout the developing CNS. Western blot analysis of whole brain lysates at E18.5 revealed a significant increase in NMIIB levels in *Mllt11* cKOs compared (Fig. 10B-C, P=.018, N=4 control, 5 cKO). We also examined Myosin 5a, NMIIA, α-tubulin, β-tubulin levels in *Mllt11* cKOs vs. control brains but did not detect any differences (data not shown). Therefore, Mllt11 interacts with NMIIB and regulates its expression in fetal mouse brains.

**Figure 10:**
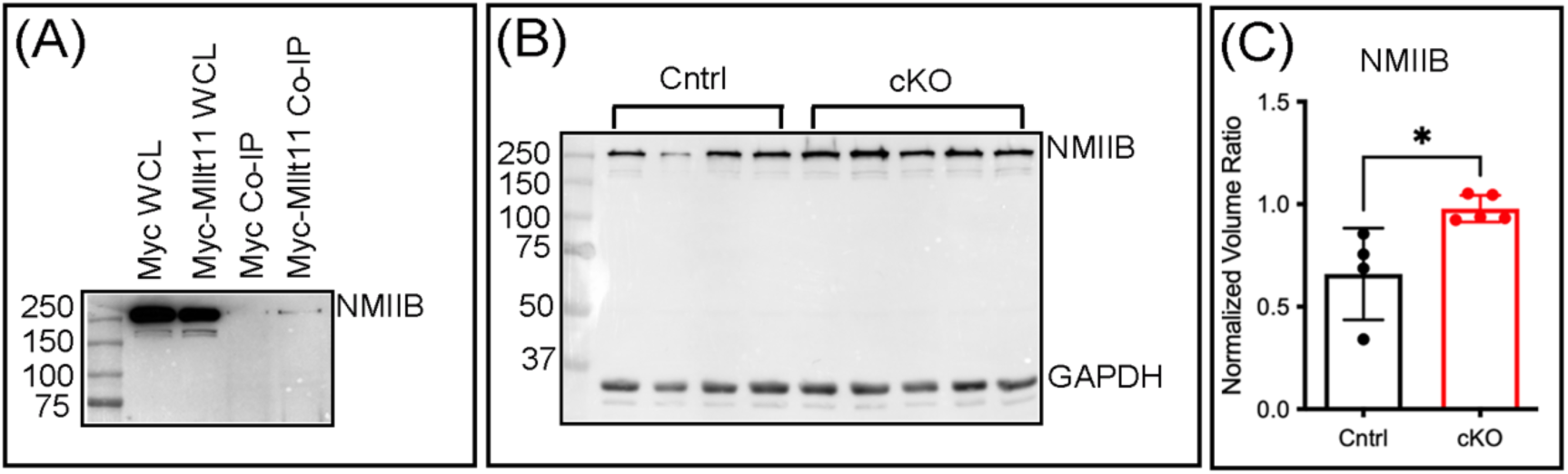
Mllt11 interacts with NMIIB. **(A)** Co-immunoprecipitation of Mllt11 and NMIIB **(B)** Western blot of NMIIB levels in whole brain lysates from E18.5 control and *Mllt11* cKO E18.5 brains. **(C)** Bar graph of comparison of NMIIB levels in control (black) versus cKO (red) western blot bands displaying a significant increase in NMIIB levels in cKO brains. Welch’s t-test, N = 5, * = p ≤ .05.

## DISCUSSION

Here we describe a role for Mllt11 in the migration of GCs during mouse cerebellar development. *Mllt11* locus activity revealed widespread expression throughout the cerebellum during embryogenesis. Through use of the *Cux2^CreERT2^* driver to target GCPs originating from the RL during early embryogenesis, we inactivated *Mllt11* specifically within the excitatory lineages of the cerebellar anlage (Capaldo & Iulianella, 2016). This approach revealed a role for Mllt11 in the formation of anchoring centres at principal fissures and both tangential and inward radial migration of GCs. In the absence of *Mllt11*, cerebella were smaller and less foliated, exhibiting shallower fissures with disrupted BG fiber patterns at their base. The reduced number of EdU^+^ cells in the dorsal lobes could explain the smaller and less developed folia observed in *Mllt11* cKOs, as folia lengthening relies on self-sustaining GC proliferation and inward migration. The aberrant BG fibers observed in *Mllt11* cKOs also likely impaired radial migration of GCs at anchoring centres and given the conditional inactivation of *Mllt11* primarily in the RL and EGL, altered glial fiber morphology was likely a result of a cell non-autonomous contribution of reduced radial invasion of GCs at folia anchor points.

The foliation defect observed in *Mllt11* mutants shares similarities, albeit to a lesser extent, with the *Engrailed 2 (En2)* mutant (Sudarov & Joyner, 2007). In *En2^-/-^* mutants the delay in anchoring centre formation leads to a smooth cerebellar surface at E17.5 instead of the presence of three principal fissures. By birth, all principal fissures have formed except the Secondary fissure, which becomes visible at P1. The altered timing of anchoring centre formation in *En2^-/-^* mutants during embryogenesis results in a pronounced postnatal phenotype, with a significantly misshapen lobule VIII (Joyner & Martin, 1987; Sudarov & Joyner, 2007). The delayed formation of the Secondary fissure in *En2^-/-^*mutants causes a reduction in fissure depth, and BG fibers fail to fan out from the anchoring centers, which resembled the *Mllt11* mutant phenotype (Sudarov & Joyner, 2007). Interesting, *En2* expression is activated by Wnt signaling and Mllt11 has previously been found to promote Wnt signaling pathways in cancer models, suggesting a possible regulatory link between the *Mllt11* and *En2* activities in the developing cerebellum (Park et al., 2015; C. O. Tse, Kim, & Park, 2017). In comparison, *Mllt11* cKOs displayed disrupted BG fibers at anchoring centres, resulting in reduced Pax6^+^ GC counts fanning out from the bases of the Preculminate and Primary fissures. These Pax6^+^ cells were likely inwardly migrating GCs which depend on the scaffolding provided by underlying BG fibers. The disrupted cerebellar foliation of *Mllt11* cKOs was also visualized by the failed invasion of Pax2^+^ interneurons into the distal columns of developing lobes. Given the that the *Cux2^CreERT)^*driver displays restricted activity to the RL, EGL and the excitatory lineages that derive from them (Capaldo & Iulianella, 2016; Iulianella et al., 2019), and not BG, the disrupted folia formation likely resulted from the effect of *Mllt11* loss on the migratory behaviour of GCs infiltrating developing cerebellar anchoring centres. This is similar to the roles we recently described for Mllt11 in regulating the migration of cortical projection neurons (Stanton-Turcotte et al., 2022) and retinal neuroblasts (Blommers et al., 2023).

Our current understanding of the mechanisms underlying cerebellar foliation is incomplete, making it challenging to interpret the disrupted GC migration as a cell intrinsic defect resulting from the movement of precursor cells from RL through the EGL, or an extrinsic defect due to altered BG scaffolds impacting radial migration from the EGL to IGL. Here we show that in the absence of *Mllt11* GCs accumulated in the RL, indicating a potential tangential migration defect. However, we also demonstrated that *Mllt11* loss was associated with fewer GCs populating the developing folia and their anchor points, reflecting altered radial migration into the IGL. This may have led to a disruption of anchoring centre cytoarchitecture and ultimately resulting in reduced foliation. These phenotypes are consistent with a role of Mllt11 in cytoskeletal regulation. Indeed, disruptions to the actin and microtubule cytoskeleton or associated motor proteins can inhibit GC migration (Bellion et al., 2005; Kawauchi & Hoshino, 2008; Trivedi & Solecki, 2011). Additionally, pharmacological inhibition of actin polymerization using cytochalasin B is sufficient to inhibit GC migration (Hatten, 1993). We recently showed that Mllt11 interacts with multiple tubulin isoforms, actin, actin-interacting proteins, and non-muscle myosins (Stanton-Turcotte et al., 2022). As the domains of Mllt11 are not yet fully defined, with no obvious cytoskeletal binding motifs, we decided to focus on confirming interactions from our proteomics list. Of these we decided to probe interactions with the non-muscle myosin NMIIB as it regulates actin and microtubule dynamics, a crucial element of growth cone steering and neuronal migration, and has been previously implicated as a regulator of neuronal migration (Ma et al., 2007; Ma et al., 2004; Rao, Hao, Wang, & Yao, 2014). We not only confirmed Mllt11 can interact with NMIIB in co-IP experiments from fetal brains, but also found that *Mllt11* loss led increased NMIIB levels, which may contribute to the altered GC migratory behaviour.

A role for NMIIB in cell motility and invasiveness has been elucidated through its ability to promote actin and microtubule dynamics by binding to and inhibiting activity of antagonistic myosin binding partner Non-muscle Myosin IIA (NMIIA) (Rao et al., 2014). It has also been implicated in regulation of cellular adhesion complexes crucial to formation of contacts between the cell and its environment, allowing for generation of frictional forces required for migration (Ma & Adelstein, 2014; Ma et al., 2007; Sahai & Marshall, 2002; Sturge, Wienke, & Isacke, 2006). Paradoxically, increased NMIIB levels have been reported in cancer cell lines, giving rise to increased motility and invasiveness (Betapudi, Licate, & Egelhoff, 2006; Ma et al., 2009). The observed increase in NMIIB levels in *Mllt11* mutant brains may therefore be due to a compensatory effort to increase cell motility due to neurogenetic defects resulting from *Mllt11* loss. Investigating this possibility will be the subject of future studies.

Cytoskeletal-associating proteins play crucial roles in cerebellar morphogenesis. For example, dystroglycan of the dystrophin-glycoprotein complex is expressed in BG endfeet and links the extracellular matrix with the intracellular actin cytoskeleton to form *glia limitans* (Moore et al., 2002). Deletion of *dystroglycan* in mice causes aberrant BG fiber organization, resulting in GC migration defects and ectopic accumulation in the EGL (Nguyen et al., 2013). Another example to highlight the integral role of the cytoskeleton comes from β-chimaerin (*Chn2*) null mice (Estep, Wong, Wong, Loui, & Riccomagno, 2018). β-chimaerin is a component of the Rho family of GTPase Activating Proteins (RhoGAPS) which is expressed in a small population of late-born premigratory GCs in the EGL and regulates other GTPases that promote actin filament depolymerization (Estep et al., 2018; Yang & Kazanietz, 2007). β-chimaerin deficiency in mice caused a subset of GCs to be arrested in the EGL, where they differentiated and formed ectopic neuronal clusters likely due to cytoskeletal dysregulation (Yang & Kazanietz, 2007). These examples highlight the critical role of the cytoskeleton in GC migration and are consistent with the view that altered cytoskeletal regulation in GCs underlie their reduced invasion at cerebellar anchoring centres in *Mllt11* cKOs, leading to reduced foliation.

In conclusion, this study identified Mllt11 as a regulator of cerebellar GC migration and foliation. We provide a possible mechanistic basis for the migratory phenotype of *Mllt11* mutant GCs by demonstrating that Mllt11 regulates NMIIB levels in the fetal mouse brain, which is a critical regulator of cellular motility and invasiveness. While this does not preclude additional roles for Mllt11 in regulating other cytoskeletal components, our findings provide additional support for its role in promoting neuronal migration during the development of laminated cytoarchitecture in the CNS.

## ACKNOWLEDGMENTS

We gratefully acknowledge funding from the Canadian Institutes of Health Research (CIHR PJT-388914), the National Science and Engineering Research Council of Canada (RGPIN 03925-20), Canadian Graduate Scholarships-Master’s award to MB, and the Killam Foundation for fellowship support to EW. We thank Sarah Whitehead for assistance with animal husbandry.

## Contributions

MB sectioned, immunostained, imaged and analyzed the data, generated figures, and wrote the first draft. DST and EAW helped with mouse embryo and fetus generation, genotyping, and manuscript preparation. MH assisted with tissue preparation. AI supervised the project, obtained funding, and edited the manuscript.

## Data availability statement

We do not have a data repository server at Dalhousie University but will make any data available upon requests.

